# Transcriptional variation in glucosinolate biosynthetic genes and inducible responses to aphid herbivory on field-grown *Arabidopsis thaliana*

**DOI:** 10.1101/563486

**Authors:** Yasuhiro Sato, Ayumi Tezuka, Makoto Kashima, Ayumi Deguchi, Rie Shimizu-Inatsugi, Misako Yamazaki, Kentaro K. Shimizu, Atsushi J. Nagano

**Affiliations:** PRESTO, Japan Science and Technology Agency, Kawaguchi 332-0012, Japan; Research Institute for Food and Agriculture, Ryukoku University, Yokotani 1-5, Seta Oe-cho, Otsu, Shiga 520-2194, Japan; Graduate School of Horticulture, Chiba University, 648 Matsudo, Chiba 271-8510, Japan; Department of Evolutionary Biology and Environmental Sciences, University of Zurich, Winterthurerstrasse 190, 8057 Zurich, Switzerland; Kihara Institute for Biological Research, Yokohama City University, 641-12 Maioka, 2440813 Totsuka-ward, Yokohama, Japan; Department of Plant Life Sciences, Faculty of Agriculture, Ryukoku University, Yokotani 15, Seta Oe-cho, Otsu, Shiga 520-2194, Japan

**Keywords:** *AOP3*, *In natura*, *Lipaphis erysimi*, RNA-Seq, Plant-insect interaction

## Abstract

Recently, increasing attempts have been made to understand how plant genes function *in natura* studies. To determine whether plant defense genes are activated under multiple biotic stimuli, we combined a high-throughput RNA-Seq with insect survey data on 19 accessions of *Arabidopsis thaliana* growing on the field site of Switzerland. We found that genes with GO annotations “glucosinolate biosynthetic process” and “response to insects” were the most significantly enriched, exhibiting largely variable expression among plant accessions. Nearly half of the total expression variation in glucosinolate biosynthetic genes, *AOPs, ESM1, ESP,* and *TGG1,* was explained by among-accession variance. Combined with the field RNA-Seq data, bioassays confirmed that *AOP3* was up-regulated in response to the mustard aphid *Lipaphis erysimi.* In addition, we also found that the expression of a major cis-jasmone activated gene *CYP81D11* was positively correlated with the number of the flea beetles *Phyllotreta* spp. The combined results from RNA-Seq and insect surveys suggested that plants can activate their defenses even when they are exposed to multiple biotic stimuli *in natura*.

## Introduction

As sessile organisms, plants are exposed to multiple stresses under naturally fluctuating environments (Wilczek et al. 2009; Carrera et al. 2017; Mishra et al. 2017). Recently, growing efforts have been made to understand how plants cope with complex field conditions (Kerwin et al. 2015; Taylor et al. 2017; Carrera et al. 2017; Kono et al. 2017; Sugiyama et al. 2017; Hiraki et al. 2018) and such *in natura* studies are important for the comprehensive understanding of gene functions from the laboratory to the field (Shimizu et al. 2011; Kudoh 2016; Yamasaki et al. 2018; Zaidem et al. 2018; Nagano et al. 2019). Insect herbivores are one of the most diverse organisms that impose biotic stresses on plants (Schoonhoven et al. 2005; Ahuja 2010; Escobar-Bravo et al. 2018). To deal with various threats, plants can activate defense mechanisms only when necessary. Such inducible defenses are triggered by wounding and insect attacks through jasmonate (JA) signaling (Mewis et al. 2005; Escobar-Bravo et al. 2017; Tsuda 2017; Zhu et al. 2018; Zhou et al. 2018; Nakano et al. 2018), while constitutive defenses are continuously expressed.

In the glucosinolate (GSL)-myrosinase system of *Arabidopsis thaliana* and related Brassicales, methionine-derived or aliphatic GSLs confer plant defenses against herbivory (Kliebenstein et al. 2002; Kerwin et al. 2015; Brachi et al. 2015) and possess natural variations in their accumulation and profiles among *A. thaliana* accessions worldwide (Kroymann et al. 2003; Chan et al. 2010; Brachi et al. 2015). The production of aliphatic GSLs is initiated by *MYB28* and *MYB29* transcription factors (Hirai et al. 2007), in which their double mutants accumulate few aliphatic GSLs (Sønderby et al. 2007). During the accumulation of aliphatic GSLs, amino acids and side chain structures are modified by methylthioalkylmalate synthase (MAM), 2-oxoglutarate-dependent dioxygenase encoded in alkenyl hydroxalkyl producing (AOP) loci, and flavin-monooxygenase glucosinolate S-oxygenase (Kliebenstein et al. 2001; Kroymann et al. 2003; Hansen et al. 2007). The enzyme myrosinase (thioglucoside glucohydrolase, TGG) catalyzes GSL and results in the emission of isothiocyanates, nitriles, or other hydrolysis products when insect herbivores bite plant tissues (Lambrix et al. 2001; Barth and Jander 2006; Zhang et al. 2006; Shirakawa and Hara-Nishimura 2018). Epithiospecifier proteins (ESP also known as TASTY: Lambrix et al. 2001; Jander et al. 2001) promote the hydrolysis of GSL with some modification by the *EPITHIOSPECIFIER MODIFIER1 (ESM1)* locus (Zhang et al. 2006), resulting in different defense activities against insect herbivores (Ratzka et al. 2002).

Insect herbivores differentially elicit defense responses of a host plant species depending on their feeding habits and host specializations. Leaf chewing herbivores crush plant tissues and accordingly, activate the GSL-myrosinase system (Barth and Jander 2006; Shirakawa and Hara-Nishimura 2018; Martinoia et al. 2018). The hydrolysis products of GSLs act as a toxin against generalist chewers (Lambrix et al. 2001; Kliebenstein et al. 2002; Barth and Jander 2006), whereas specialist herbivores exploit GSL and its hydrolysis products as a host plant signal (Ratzka et al. 2002; Renwick et al. 2006). Sapsuckers, such as aphids and thrips, consume plant fluids and, very rarely, crushes plant tissues (Mewis et al. 2005; Kempema et al. 2007). In addition, damaged plants may emit volatile chemicals that elicit defenses of other individual plants or alter feeding behaviors of other insect species (Bruce et al. 2008; Matthes et al. 2011; Yazaki et al. 2017). To date, *A. thaliana* is known to possess natural variations in secondary metabolisms and inducible defenses (Chan et al. 2010; Snoeren et al. 2010; Routaboul et al. 2012); however, how they function in nature remains poorly understood.

In wild populations, *A. thaliana* has multiple life-cycles within a calendar year (Thompson 1994; Wilczek et al. 2009; Taylor et al. 2017) and is attacked by various herbivores (Arany et al. 2008; Harvey et al. 2008; Sato et al. 2018). Sato et al. (2018) observed both flowering and vegetative *A. thaliana* co-occurring and being wounded by insect herbivores in a wild population near Zurich, Switzerland in summer. By simulating the summer cohort, our previous study found that 12 insect species, including mustard aphids *Lipaphis erysimi* (Homoptera), flea beetles *Phyllotreta striolata* and P *atra.* (Coleoptera), diamondback moths *Plutella xylostella* (Lepidoptera), and the western flower thrips *Frankliniella occidentalis* (Thysanoptera), colonized *A. thaliana* and such an insect community composition significantly varied among *A. thaliana* accessions (Sato et al. 2018). We also found that plant trichomes play a key role in altering the abundance of leaf chewing herbivores, but there were unclear correlations between insect abundance and laboratory-measured GSL profiles (Sato et al. 2018). To link insect abundance and plant physiological status in the field, we employed transcriptomics, which were widely used to reveal comprehensive pictures of gene expression (Sun et al. 2017; Xu et al. 2018; Wang et al. 2018; Lin et al. 2018; Want et al. 2018). Here we adopted our previously established protocol of cost-effective RNA-Seq (Nagano et al. 2015; Kamitani et al. 2016; Ishikawa et al. 2017) to the field-grown *A. thaliana.*

The purpose of this study was to reveal to what extent variations in gene expression could be explained by plant genotypes under field conditions, and then to specify which herbivores could modulate plant defense responses. To address these issues, we combined a high-throughput RNA-Seq with insect monitoring data on 19 accessions of *A. thaliana* individuals (Table 1; Figure 1). Such a joint approach using the field transcriptome analysis and insect surveys will provide an overall picture of how *A. thaliana* responds to multiple attackers under natural field conditions.Insect herbivores on field-grown *A. thaliana*

**Figure 1.**
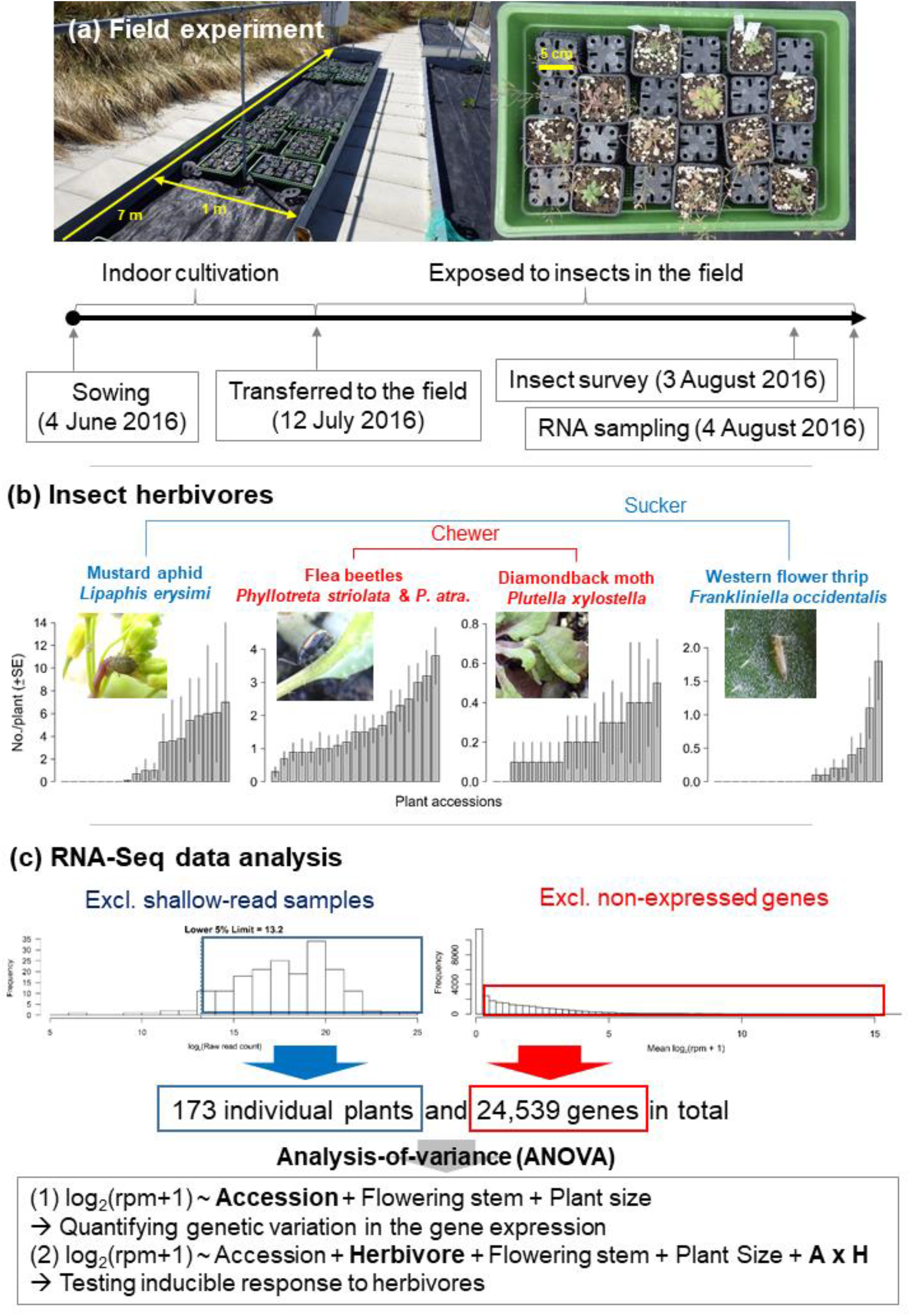
Outline of this field study on *Arabidopsis thaliana.* (a) Procedure of the field experiment. (b) Observed variation in insect abundance among plant accessions. (c) Filtering and statistical analysis of RNA-Seq data. In the formula of ANOVA, the Herbivore represents the main effect of the number of herbivores while the A × H indicates the interaction term between the plant accession and the number of herbivores.

**Table 1.**
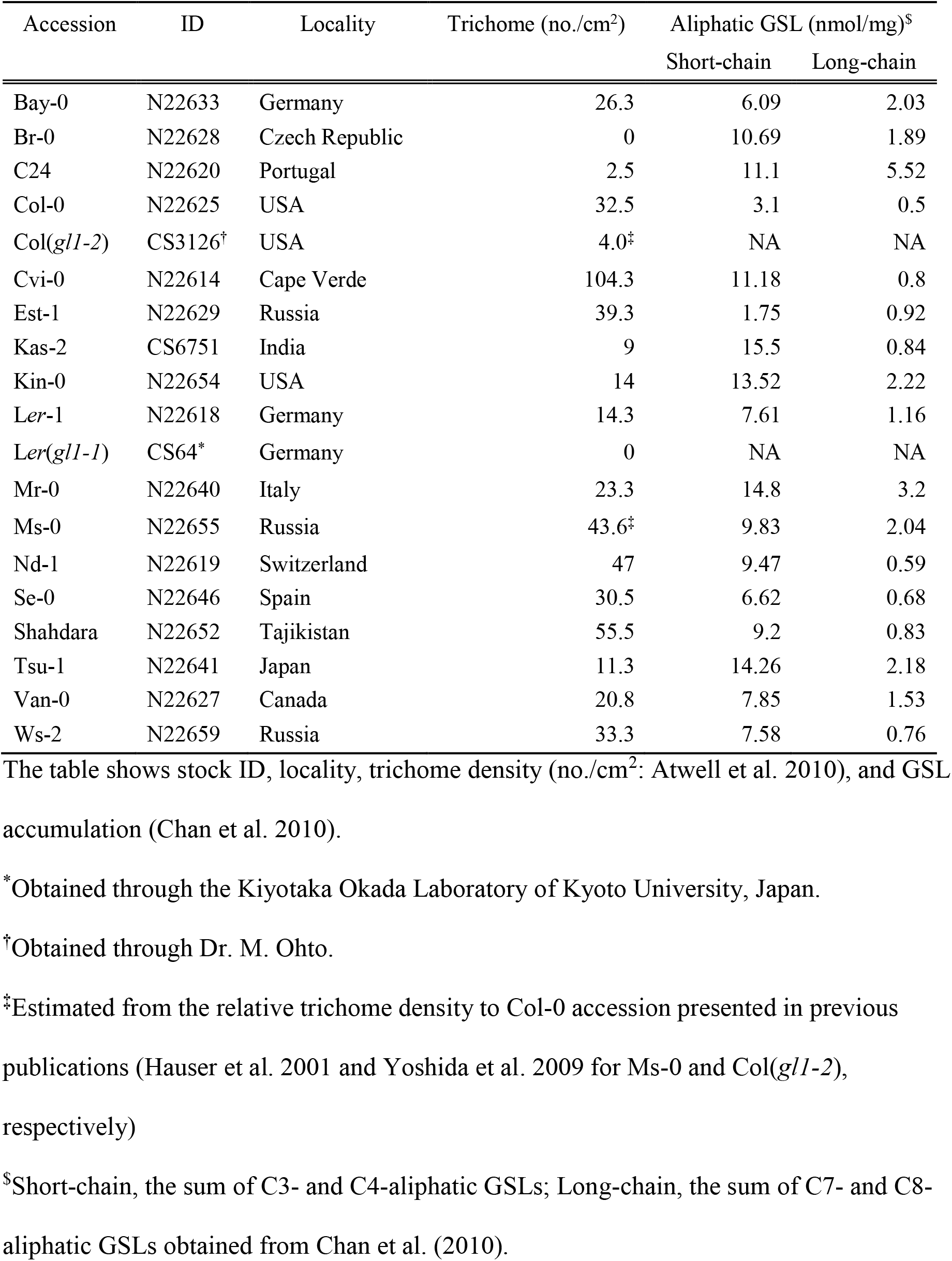
List of *Arabidopsis thaliana* accessions used in this study.

## Results

### *Insect herbivores field-grown* A. thaliana

We exposed a summer cohort of *A. thaliana* to the field environment in Zurich, Switzerland (47°23’N, 8°33’E) (Figure 1a). As major insect herbivores, we observed *L. erysimi, P. striolata, P. atra., P. xylostella,* and *F. occidentalis*. Of these herbivores *L. erysimi* and *F. occidentalis* are sucking insects, while *P. xylostella* and *Phyllotreta* are leaf chewers (Ahuja 2010; Escobar-Bravo et al. 2018). *Lipaphis erysimi, P. xylostella,* and *Phyllotreta* are specialists of Brassicaceae (Ahuja 2010), while *F. occidentalis* is a generalist, which feeds on various plant families (Escobar-Bravo et al. 2018) (Figure 1b).

### *Gene expression variation among* A. thaliana *accessions*

Leaf samples for the RNA-Seq analysis were collected from 183 individual plants. Given that previous laboratory experiments detected inducible defenses 24-48 h after insect attacks (e.g., Mewis et al. 2005; Kuśnierczyk et al. 2008; Matthes et al. 2011), the leaf sampling for RNA-Seq was done 1 d after the final insect monitoring (Figure 1a). We sequenced 92 samples per lane using Illumina HiSeq^®^ 2500 and obtained 829,681 mapped reads per sample on average. After discarding the lower 5% shallow-read samples and excluding genes whose expression level was zero in mean log_2_ (rpm + 1) (Figure 1c), 173 individual plants with 24,539 genes were subject to our statistical analyses.

A type III Analysis-of-variance (ANOVA: Sokal and Rolf 2012) was performed to partition whole expression variation into the effects of the plant accession, initial size of individual plants, the presence of a flowering stem, and unexplained residuals (Figure 1c). This variation partitioning showed that, when ordered by its expression variation explainedby each factor, the top 5% of the genes had more than 20% variation attributable to plant accessions (Figure 2a; Table S1). In these highly variable genes, 22 gene ontologies (GO) were significantly enriched at *P*_FDR_ < 0.05 (Table S2). Gene ontology annotation of “response to insect” and “glucosinolate biosynthetic process” were most and second most significantly enriched, respectively (Table 2).

**Figure 2.**
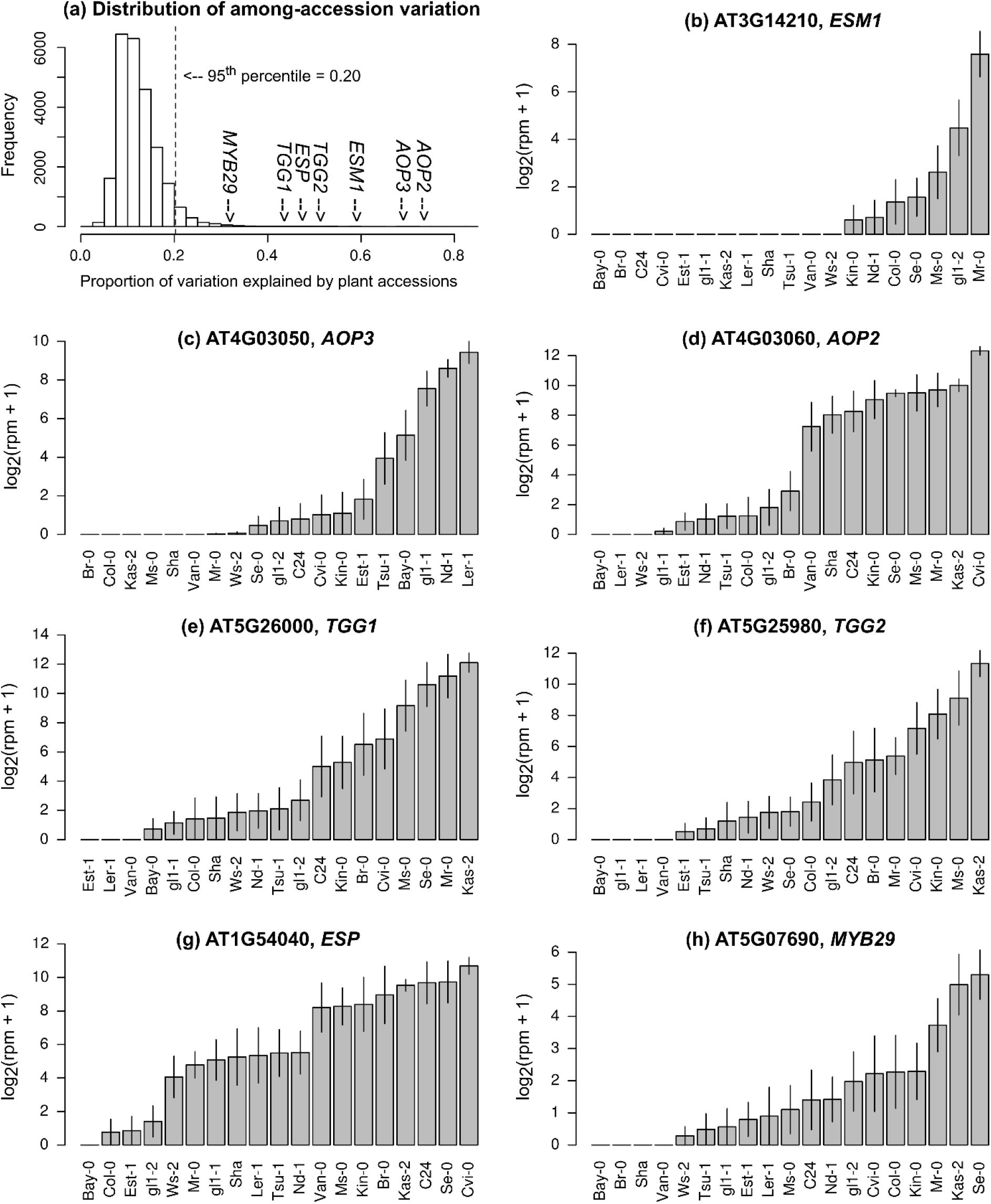
Natural variation in the expression levels of genes involved in glucosinolate biosynthesis and hydrolysis. (a) Histogram showing the proportion of variation explained by plant accessions, (b) the expression of *ESM1,* (c) *AOP3,* (d) *AOP2,* (e) *TGG1,* (f) *TGG2,* (g) *ESP,* and (h) *MYB29.* Grey bars and vertical lines indicate mean ± SE. The list of the top 5% variable genes is available in supporting information (Table S1).

**Table 2.** Gene Ontology (GO) enrichment analysis of genes exhibiting the top 5% expression variation explained by plant accessions. Shown are the top five significant GOs within the biological processes. P-values are corrected by the false discovery rate (*P*_FDR_). The entire result at *P*_FDR_ <0.05 is available in Table S2.

| GO ID | Term | $P_{\text{FDR}}$ |
| --- | --- | --- |
| GO:0009625 | response to insect | 4.76E-05 |
| GO:0019761 | glucosinolate biosynthetic process | 6.35E-05 |
| GO:0055114 | oxidation-reduction process | 0.000116 |
| GO:0009627 | systemic acquired resistance | 0.000327 |
| GO:0009414 | response to water deprivation | 0.000446 |

Several key genes of aliphatic GSL biosynthesis, such as *AOPs* (Kliebenstein et al. 2001), showed over half of the expression variation attributable to plant accession (Figure 2a). *AOP2* and *AOP3* are tandemly located in the genome and encode 2-oxoglutarate-dependent dioxygenases that are involved in the side-chain modification of aliphatic GSLs (Kliebenstein et al. 2001). Genes involved in GSL hydrolysis, such as *TGGs, ESM1,* and *ESP1* (Lambrix et al. 2001; Zhang et al. 2006; Barth and Jander 2006), also exhibited such a large variation in expression that these genes had nearly a half variation attributable to the plant accession (Figure 2b, e, f, g). *TGG1* and *TGG2* are functionally redundant and their double mutants are known to be susceptible to generalist caterpillars, but not to aphids and specialist caterpillars (Barth and Jander 2006). *ESM1* and *ESP* were initially screened by QTL mapping utilizing the natural variation between the Col and L*er* accession (Lambrix et al. 2001; Jander et al. 2001; Zhang et al. 2006). Furthermore, the transcription factor gene *MYB29,* which is responsible for the high accumulation of aliphatic GSLs (Hirai et al. 2007) showed 32% variation in its expression among field-grown accessions (Figure 2 h).

### Inducible response to leaf chewing and sapsucking herbivores

To find candidate genes possessing inducible responses, we included the effects of the number of each herbivore into the ANOVA (Figure 1c). In the ANOVA, the main effect of the number of herbivores (Figure 1c) was used to determine whether the gene expression level was correlated with herbivore abundance, while the interaction term of the number of herbivores with plant accessions (A×H, Figure 1c) addressed whether the magnitude of the correlation differed among plant accessions. The ANOVA found 27, 25, 20, and 19 candidate genes were significantly related to inducible responses to the mustard aphids, the flea beetles, the diamondback moths, and the western flower thrip, respectively (Table S3).

The expression of *AOP3* was significantly correlated with the number of mustard aphids, *L. erysimi* (*P*_FDR_ < 0.001: Figure 3a, Table S3), indicating that its induction by the aphids depended on background genomic variation. This explained the additional 7% variation in *AOP3* expression. Similar interactions with aphid herbivory were observed for *MYB113* and *JAX1.* Furthermore, *MYB113* is involved in anthocyanin biosynthesis and induced via JA signaling (Gonzalez et al. 2011), and *JAX1* encodes jacalin-type lectin resistance to potexvirus and exhibits varying resistance among natural accessions (Yamaji et al. 2012).

**Figure 3.**
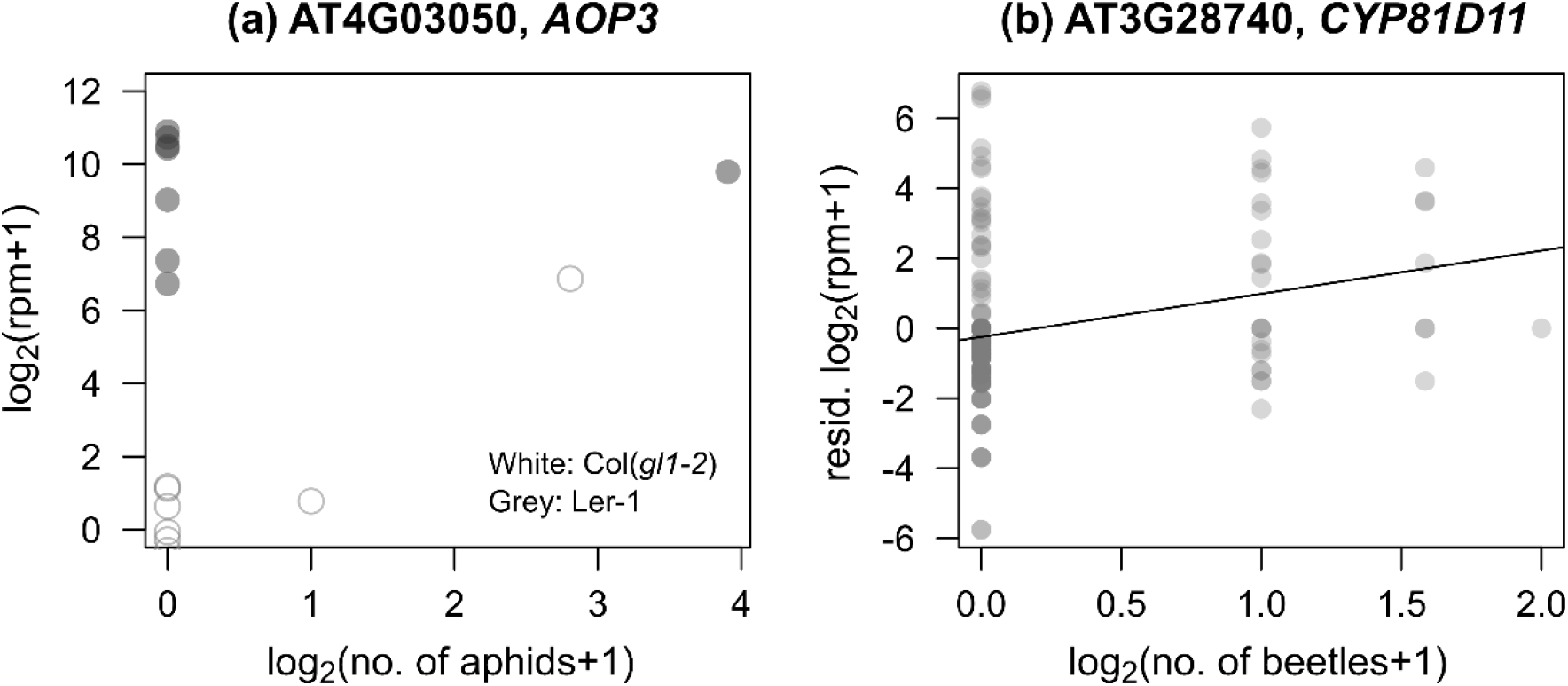
Candidate genes showing a positive relationship between their expressions and insect abundance in the field. (a) Relationship between the number of specialist mustard aphids *Lipaphis erysimi* and the expression of *AOP3* in a constitutively expressed accession Ler-1 or a potentially induced accession Col(gl1-2). (b) Relationship between the number of *Phyllotreta* beetles and the expression of *CYP81D11.* Residuals of log_2_(rpm + 1) of *CYP81D11* normalized the expression difference among plant accessions.

In the same manner, the number of flea beetles *(Phyllotreta* spp.) was positively correlated with the expression level of a major cis-jasmone activated gene, *CYP81D11* (Matthes et al. 2010, 2011), (*P*_FDR_ = 0.027: Figure 3b, Table S3). However, its expression was not correlated with plant accessions (plant accession × beetles, *P*_FDR_ = 0.99), indicating limited effects of background genomic variation. Additionally, the number of leaf holes made by the flea beetles was only related to the expression of three loci of unknown function, AT2G41590 (plant accession × holes, *P*_FDR_ < 10^-6^), AT1G34844 (plant accession × holes, *P*_FDR_ < 10^-13^), and AT2G47570 (plant accession × holes, *P*_FDR_ = 0.007).

In addition, AT5G48770, which encodes disease resistance proteins of the TIR-NBS-LRR class family and has GO annotations of “defense response” was expressed in response to the diamondback moth P *xylostella* (Table S3).

Finally, the presence of the western flower thrips, *F. occidentalis,* resulted in the expression of one locus, AT2G15130, that encodes a plant basic secretory protein family protein and has the GO annotation “defense response” (Table S3).

### Laboratory bioassay using the specialist aphids

Based on the statistical interactions between plant accession and the number of aphids, we found a positive correlation between *AOP3* and the number of the mustard aphids, *L. erysimi,* in a plant with the Col genomic background (Figure 3a). To determine the possibility of an inducible response of *AOP3* to aphid herbivory, we released *L. erysimi* on Col-0 under controlled conditions and quantified the expression of *AOP3* with RT-qPCR (Figure 4a, b). The bioassay confirmed that the expression of *AOP3* was up-regulated in aphid-infested plants (Wilcoxon rank sum test, *W* = 0, *n* = 16, *P* = 0.0002: Figure 4c).

**Figure 4.**
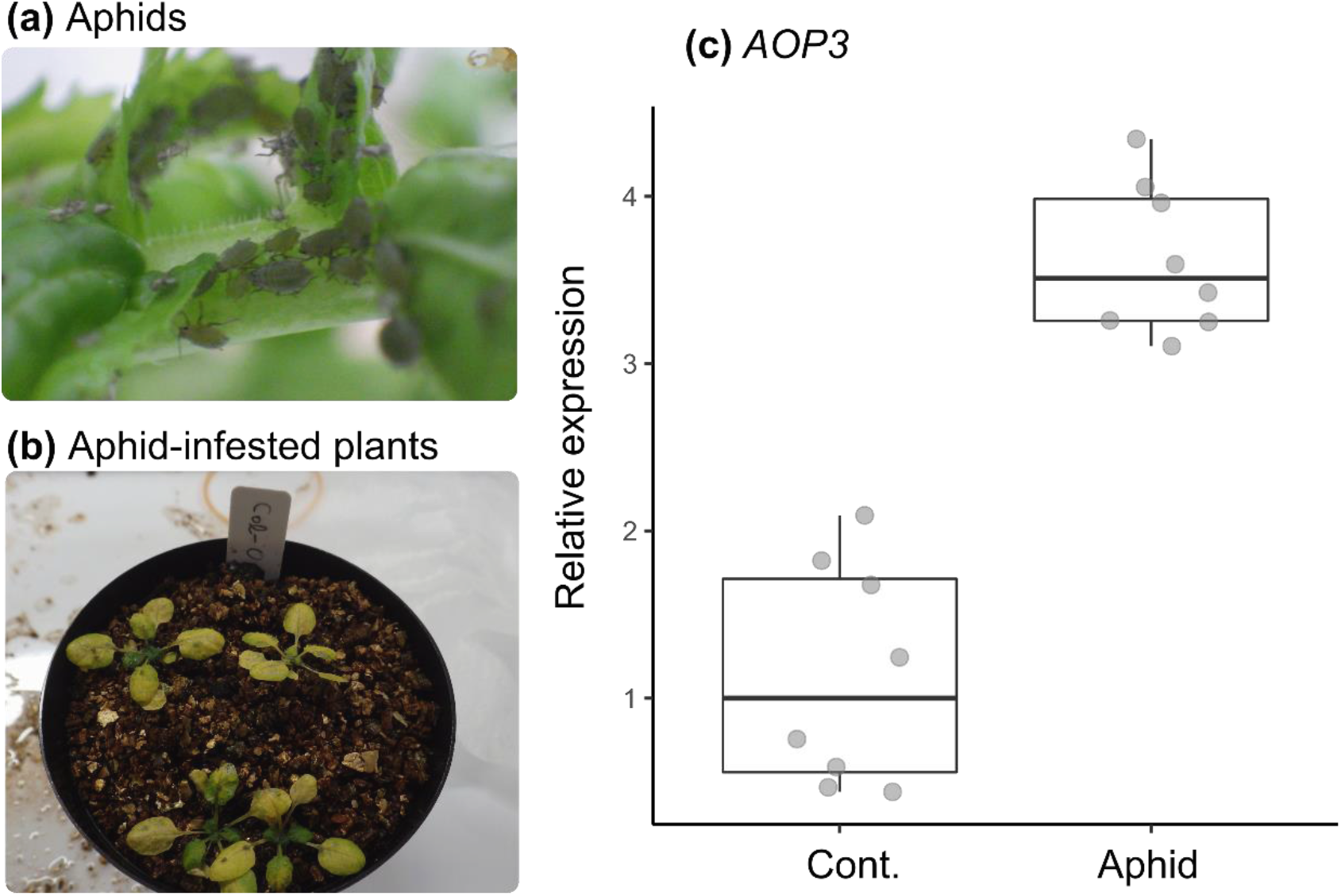
Inducible response of *AOP3* to the specialist mustard aphid *Lipaphis erysimi* in the laboratory-grown Col-0 accession. (a) Photograph of laboratory-reared colony of *L. erysimi.* (b) Aphid-infested seedlings of Col-0 accession. (c) The RT-qPCR analysis of *AOP3* in aphid-infested and control plants. Seedlings are infested by aphids for seven days (Aphid), and no aphids were released for the controls (Cont.).

## Discussion

### Expression variation in glucosinolate biosynthetic genes

Glucosinolate profiles, leaf damage, and plant fitness significantly varied between the GSL wildtype and mutants as shown by a field study on *A. thaliana* (Kerwin et al. 2015). Despite the fact that insect abundance and other environmental conditions were not manipulated, our study detected the GO enrichment of “GSL biosynthesis” and “response to insects” in genes showing the top 5% variation among *A. thaliana* accessions (Table 2). Notably, nearly half of the expression variation in *AOPs, ESM,* and *TGG1,* was explained by plant accessions (Figure 2), showing a comparable magnitude of variation with the heritability reported by a laboratory eQTL study (Wentzell et al. 2007). A third of the expression variation in the transcription factor gene, *MYB29,* was attributable to plant accessions, even though this gene is known to respond to water stress and other abiotic stimuli (Mewis et al. 2012; Martínez-Ballesta et al. 2015; Zhang et al. 2017). Overall, our genome-wide analysis using RNA-Seq indicated that GSL biosynthesis and its anti-herbivore functions were one of the most genetically variable functions in field-grown *A. thaliana.*

Among the GSL biosynthetic genes, *AOPs* showed a remarkably large expression variation among natural accessions. More specifically, Ler-1 accession expressed *AOP3* and not *AOP2,* while Cvi expressed *AOP2,* but not *AOP3* (Figure 2c, 2d) as reported by Kliebenstein et al. (2001). In the Col accession, *AOP2* encodes non-functional proteins (Kliebenstein et al. 2001), whereas, *AOP3* is not expressed in leaves without herbivory (Schmid et al. 2005). Strong genome-wide associations between the *AOP* loci and GSL profiles have been repeatedly detected among natural accessions cultivated under laboratory (Chan et al. 2010) or controlled greenhouse (Brachi et al. 2015) conditions. Based on a genome scan, Brachi et al. (2015) also detected an adaptive differentiation in the *AOP* loci within the European *A. thaliana.* In this evolutionary context, this study provided field evidence for a link between genomic and functional variations in *AOPs.*

### Genes possessing inducible responses to herbivory

Consistent with field RNA-Seq data, our laboratory bioassay revealed that the Col-0 accession had an inducible response in *AOP3* to the mustard aphid *L. erysimi.* This aphid species as well as the generalist and cabbage aphids, *Myzus persicae* and *Brevicoryne brassicae,* are major natural enemies of *A. thaliana* (Züst et al. 2012). In the ecological context, previous studies reported unclear geographical associations between the three aphid species and AOP-related chemotypes in Europe, though the geographical distribution of the aphids was linked to that of MAM-related chemotypes (Züst et al. 2012). Our previous study also reported no significant correlations between laboratory-measured profiles of aliphatic GSL and the abundance of *L. erysimi* in the field. Microarray analyses and the results of this study, showed that the aphid species differentially induced *AOP3* as this gene could be up-regulated in Col-0 by *L. erysimi,* down-regulated in Ler by *B. brassicae* (Kuśnierczyk et al. 2008), and not induced in Col by *M. persicae* (Kempema et al. 2007). Given its natural variation and response to different aphid species, the inducible response of *AOPs* might be a clue for understanding why *AOP* loci are not tightly linked to the higher phenotypes, such as aphid resistance, in wild populations.

Of the genes with the GO annotation of “response to insects”, *CYP81D11* exhibited a significantly positive correlation between its expression and the abundance of flea beetles. Previous studies revealed that *CYP81D11* was up-regulated by multiple biotic stimuli, including insect herbivory and pathogen infection (Matthes et al. 2011) via cis-jasmone, a plant volatile emitted by wounding (Bruce et al. 2008; Matthes et al. 2010; Matthes et al. 2011). While Matthes et al. (2011) used a Col background as a standard accession, this study included multiple natural accessions and found a positive correlation between flea beetle abundance and *CYP81D11* expression (Figure 3b). However, there was no significant correlation between *CYP81D11* expression and the number of leaf holes made by these beetles. This result was probably because the leaf holes remained on the leaves for a few weeks and did not reflect the timing of wounding. The results of *CYP81D11* indicated that not only herbivory, but also the insect abundance data, were needed to detect its inducible response to flea beetles, exemplifying the importance of detailed ecological observations in *in natura* studies.

### Conclusion

The combination of insect surveys and field transcriptome analyses led us to observe an inducible defense against insect herbivores on *Arabidopsis thaliana*. These results provide field evidence that the molecular machinery of *Arabidopsis* defense can function in noisy environments. While previous field studies on a Brassicaceae crop reported significant transcriptional changes in response to entire herbivore communities (Broekgaarden et al. 2010), our large-scale RNA-Seq and insect monitoring dissected such a transcriptional response to each herbivore. Since the insect species studied here are also known as worldwide herbivores feeding on cultivated and wild Brassicaceae (Yano 1994; Ahuja et al. 2010; Sato and Kudoh 2017), our findings may provide molecular insights into Brassicaceae-herbivore interactions *in natura*.

## Materials and Methods

### Field experiment

In the field experiment, we used 17 natural accessions and two glabrous mutants (Table 1), which covers phenotypic variation in trichomes (Larkin et al. 1999; Atwell et al. 2010; Bloomer et al. 2012) and glucosinolates (Chan et al. 2010). We initially prepared 10 replicates of the 19 accessions (= 190 plants in total) in an environmental chamber, and then transferred them to the outdoor garden of the University of Zurich at Irchel campus (Zurich, Switzerland: 47°23’N, 8°33’E, alt. ca. 500 m) (Figure 1a). Plants were cultivated using mixed soils of agricultural composts (Profi Substrat Classic CL ED73, Einheitserde Co.) and perlites with a compost to perlite ratio of 3:1 litre volume. No additional fertilizers were supplied because the agricultural soils contained fertilizers. Seeds were sown on the soil and stratified under constant dark conditions at an ambient temperature of 4 °C for a week. Plants were grown under short-day conditions (8: 16 h light: dark [L:D] at 20 °C and a relative humidity of 60%) for a month. The tray positions were rotated every week to minimize the growth bias based on light conditions. Individual plants were moved to a plastic pot (6.0 × 6.0 × 6.0 cm) and acclimated for 3 days in a shaded area outdoors prior to field experiments. The potted plants were randomly placed in a checkered manner between three blocks where 68, 69, and 53 plants were assigned within each block. The potted plants were set on water-permeable plastic sheets without being embedded in the ground (Figure 1a). Each block was more than 1.0 m apart from the other and the plants were watered every morning and dawn. These experiments were conducted from July 13 to August 3, 2016.

Insects on individual plants were visually counted every 2-3 days. We counted these herbivores and the leaf holes made by the flea beetles. The initial plant size (evaluated by the length of the largest leaf at the start of field experiment: mm) and presence of flowering stems (two weeks after the start of experiment: binary variable) were also recorded so we could consider these phenotypes as covariates of statistical analyses. All monitoring was conducted by a single observer during the daytime (08:00-17:00) for 3 weeks after the beginning of the field experiment to minimize variation. Details of insect abundance and diversity are reported in our previous publication (Sato et al. 2018). In case wounding activated plant defense responses via JA signaling (e.g., Mewis et al. 2005; Broekgaarden et al. 2010; Matthes et al. 2010), we did not sample any leaves until the end of the field experiment.

### RNA-Seq experiments and data filtering

Leaves were collected from field-grown plants at the end of the experiment (August 4, 2016). The leaf samples were immediately soaked in RNA preservation buffer (5.3 M (NH_4_)_2_SO_4_, 20 mM EDTA, 25 mM Trisodium citrate dihydrate, pH 5.2) at 4 °C overnight and stored at −80 °C until RNA extraction. Total RNA was extracted using the Maxwell 16 Lev Plant RNA Kit (Promega) according to the manufacturer’s protocol. Selective depletion of rRNAs and highly abundant transcripts were conducted prior to RNA-Seq library preparations as previously described (Nagano et al. 2015). Then, RNA-Seq library preparation was performed as previously described (Ishikawa et al. 2017). Sequencing using Illumina HiSeq^®^ 2500 was carried out by Macrogen Co.

The fastq files generated by the sequencing were preprocessed using trimmomatic version 0.32 (Bolger et al. 2014). The preprocessed sequences were mapped on the *A. thaliana* reference genome (TAIR10 cDNA) using bowtie version 1.1.1 (Langmead et al. 2009) and then quantified using RSEM version 1.2.21 (Li and Dewey 2011). The parameter setting of trimmomatic, bowtie, and RSEM were the same as described by Kamitani et al. (2016). According to Kamitani et al. (2016), we calculated the raw read count and read per million (rpm) from the expected read count generated with RSEM. Transposable elements were excluded prior to statistical analyses. We calculated the total raw read counts for each plant sample and discarded shallow-read samples belonging to the lower 5th percentile of the total raw read counts (Figure 1c). Consequently, samples with more than 12,130 reads were subject to statistical analyses. To exclude non-expressed genes, we then averaged log_2_ (rpm + 1) for each gene between all plant samples and eliminated genes with a zero average of log_2_(rpm + 1) (Figure 1c). Overall, we obtained the final dataset on 24,539 genes for 173 plants. In this final dataset, 53 out of 173 samples had < 10^5^ total reads. However, overall trends did not change when we set the threshold at 10^5^, although its statistical power decreased due to the sample size limitation.

### Statistical analysis

We used a Type III analysis-of-variance (ANOVA: Sokal and Rolf 2012) to screen genes showing a large expression variation among accessions (Figure 1c). We formulated the liner model as: Y ~ Accession ID (factorial) + Flowering stem (0/1) + Initial leaf length (mm), where Y indicates log_2_(rpm + 1) of a focal gene. Sum-of-squares (SS) were calculated to partition expression variation attributable to each explanatory variable. The proportion of expression variation explained by the plant accession was evaluated as SS of the plant accession ID divided by the total SS. Genes in the top 5% of expression variation were selected and subject to statistical analysis, as described below. All statistical analyses were performed using R version 3.2.0 (R Core Team 2015).

Subsequently, gene ontology (GO) enrichment analysis was applied to genes showing the top 5% values of the proportion of expression variation explained by the plant accession. The GO.db package (Carlson 2017) and TAIR10 gene annotation were used to build the in-house R script of GO enrichment analysis. The statistical significance of the GO term was determined using Fisher’s exact probability tests against the entire database. The p-values were adjusted by the false discovery rate (FDR: Benjamini and Hochberg 1995) using the p.adjust function of R. When significant GO terms were detected, the GOBPOFFSPRING database in the GO.db package was used to find the most descendant GO within the Biological Process.

We then incorporated the effects of herbivores into an ANOVA to determine whether insect herbivory altered gene expression among plant accessions. We formulated the linear model in ANOVA as: Y ~ Accession ID (factorial) + No. of herbivores + Flowering stem (0/1) + Initial leaf length (mm) + (Accession ID × No. of herbivores), where no. of herbivores referred to the number of individuals of a focal herbivore species. The interaction term, Accession ID × No. of herbivores, represented the non-additive, combined effect exerted by the plant accession and the number of herbivores. This ANOVA was repeated for four major herbivores: The number of mustard aphids *Lipaphis erysimi, Phyllotreta* beetles, leaf holes made by *Phyllotreta* spp., diamondback moths *Plutella xylostella,* and western flower thrips *Frankliniella occidentalis.* The number of herbivores was log-transformed to improve normality. Given that previous laboratory experiments detected inducible defenses 24-48 h after insect attacks (e.g., Mewis et al. 2005; Kuśnierczyk et al. 2008; Matthes et al. 2011), we used insect abundance data on August 3, 2016 i.e., a day before RNA sampling as an explanatory variable. The *p*-values were calculated using the *F*-test and corrected by FDR. The aov function implemented in R was used to perform the ANOVA, with each factor dropped from the full model.

### Laboratory bioassay and RT-qPCR

To determine whether *A. thaliana* possess inducible responses to the mustard aphid, *L. erysimi*, we released this aphid species on Col-0 accessions under controlled conditions in the laboratory, and then quantified the expression of *AOP3* gene in intact and infested plants (Figure 4). *Lipaphis erysimi* were collected from *Rorippa indica* growing in the Seta campus of Ryukoku University, Japan (34°58’N, 135°56’E), and maintained on leaves of *Raphanus sativus* var. *longipinnatus* before the bioassay. Seeds were sown in plastic pots (6 cm in diameter and height) filled with moist vermiculate. Four seedlings were kept per pot and others were discarded after germination. Seedlings were grown under 16L: 8D conditions at an ambient temperature of 20 °C for a month. Liquid fertilizer was diluted 2000-times diluted and supplied during the cultivation (Hyponex, Hyponex Japan, Osaka; N:P:K = 6:10:5). We assigned two pots to the aphid treatment, while the other two pots were the controls. Approximately 80 wingless aphids were released per pot for the aphid treatment, and the pots were separately covered with an unwoven net. Leaf sampling was conducted once a week after the release of aphids. Leaves were soaked in RNA preservation buffer (5.3 M (NH_4_)_2_SO_4_, 20 mM EDTA, 25 mM Trisodium citrate dihydrate, pH 5.2) overnight and stored at −80 °C until RNA extraction.

Total RNA was extracted using a Maxwell 16 Lev Plant RNA Kit (Promega, Tokyo, Japan) and RNA concentration was measured using a Quant-iT RNA Assay Kit Broad Range (Invitrogen, Carlsbad, CA, USA). cDNA was synthesized from 300 ng of the total RNA using the reaction solution composed of 10 μl of template RNA, 4.0 μl of 5× SuperScript IV Reverse Transcriptase buffer (Invitrogen, Carlsbad, CA), 0.5 μl of RNasin^®^ Plus RNase inhibitor (Promega, Tokyo, Japan), 2.0 μl of 100 mM DTT (Invitrogen, Carlsbad, CA), 0.4 μl of 25 mM each dNTP (Clontech, Palo Alto, CA), 0.5 μl of SuperScript IV Reverse Transcriptase (Invitrogen, Carlsbad, CA, USA), and 0.6 μl of 100 μM random primer (N)6 (TaKaRa, Kusatsu, Japan), with RNase-free water added up to 20 μl. For the reverse transcription step, the mixture was incubated at 25 °C for 10 min, followed by 50 min at 56 °C. SuperScript IV was inactivated by heating the mixture at 75 °C for 15 min. The cDNA was 10× diluted with RNase-free water and then RT-qPCR was performed using the Roche LightCycler^®^ 480 with 10 μl reaction solution composed of 2 μl of the template, 5 μl of KAPA SYBR Fast RT-qPCR solution (Kapa Biosystems, Inc., Woburn, MA), 0.5 μl of 10 μM forward and reverse primers, and 2 μl of RNase-free water. The forward and reverse primer of the target gene *AOP3* was 5’-TCAGGGGTCGGTTTTGAAGG-3’ and 5^1^-GTGAAAGGTTTCGGGCACAC-3^1^, respectively. *ACT2* and *EF-1α* were measured for the internal controls (see Czechowski et al. 2005 for primer sequence). The cDNAs were amplified following denaturation, using the 35-cycle programs (10 s at 95 °C; 20 s at 63 °C;10 s at 72 °C per cycle). Three technical replicates were set for individual plants and primers. The primer of the target gene *AOP3* was designed based on its full length CDS using NCBI Primer-BLAST with a product length parameter of 50-150 bp. We tried six candidate primers and selected the *AOP3* primer above based on the melting curve of laboratory-grown Col-0 and Ler-1 accessions. Cp values were calculated following the second derivative maximum method and averaged among three technical replicates. The geometric mean of Cp values between *ACT2* and *EF-1α* was used as the internal control. Delta Cp values were calculated for each individual plant between the target and internal control. Wilcoxon rank sum test was used to test differences in the delta Cp values between the intact and aphid-infested plants.

## Acknowledgements

The authors thank F. Kobayashi for assistance in RNA extraction, and Dynacom Co. Ltd. for help with the RNA-Seq data analyses.

## Supporting information

**Table S1.** List of genes showing the top 5% expression variation among *Arabidopsis thaliana* accessions.

[TableS1_VarTop5% sheet]

**Table S2.** Gene ontology (GO) categories enriched at *P*_FDR_ < 0.05 in genes showing the top 5% variation explained by plant accessions. *P*-values were corrected by the false discovery rate (*P*_FDR_).

[Table S2_GO sheet]

**Table S3.** List of candidate genes possessing inducible response to sapsucking and leaf chewing herbivores at *P*_FDR_ < 0.05. (a) The mustard aphid *Lipaphis erysimi,* (b) the flea beetles *Phyllotreta* spp., (c) the diamondback moth *Plutella xylostella,* and (d) the western flower thrip *Frankliniella occidentalis.* NA means not available. P-values were corrected by the false discovery rate (*P*_FDR_). Candidate genes are listed for the main effect of each herbivore (Main) and its interactive effect with plant accessions (Interaction).

[Table S3_GxE sheet]

## Abbreviations

ANOVA: analysis of variance;
AOP: alkenyl hydroxalkyl producing;
ESM1: epithiospecifier modifier 1;
ESP: epithiospecifier protein;
FDR: false discovery rate;
GO: gene ontology;
GSL: glucosinolate;
MAM: methylthioalkylmalate synthase;
rpm: read per million;
TGG: thioglucoside glucohydrolase

